# Ecological drivers of jellyfish blooms – the complex life history of a ‘well-known’ medusa (*Aurelia aurita*)

**DOI:** 10.1101/102814

**Authors:** J. Goldstein, U.K. Steiner

## Abstract

1. Jellyfish blooms are conspicuous demographic events with significant ecological and socio-economic impact. Despite worldwide concern about an increased frequency and intensity of such mass occurrences, predicting their booms and busts remains challenging.
2. Forecasting how jellyfish populations may respond to environmental change requires taking into account their complex life histories. Metagenic life cycles, which include a benthic polyp stage, can boost jellyfish mass occurrences via asexual recruitment of pelagic medusae.
3. Here we present stage-structured matrix population models with monthly, individual-based demographic rates of all life stages of the moon jellyfish *Aurelia aurita* (sensu stricto). We investigate the life stage-dynamics of these complex populations under low and high food conditions to illustrate how changes in medusa density depend on non-medusa stage dynamics.
4. We show that increased food availability can be an important ecological driver of jellyfish mass occurrences, as it can temporarily shift the population structure from polyp- to medusa-dominated. Projecting populations for a winter warming scenario additionally enhanced the booms and busts of jellyfish blooms.
5. We identify demographic key variables that control the intensity and frequency of jellyfish blooms in response to environmental drivers such as habitat eutrophication and climate change. By contributing to an improved understanding of mass occurrence phenomena, our findings provide perspective for future management of ecosystem health.

## Introduction

The biodiversity, productivity and resilience of marine ecosystems worldwide are threatened by an imbalance in the ecological parameters regulating population outbreaks. Mass occurrences of micro- and macroalgae, cnidarians, ctenophores, molluscs, echinoderms, tunicates and fish appear with increasing frequency and intensity at a global scale in recent decades (Breithaupt, 2003; Lucas & Dawson, 2014; Maji, Bhattacharyya, & Pal 2016). Despite mass occurrences and mass mortality events are widespread – and can provide major implications for marine ecosystem health – predictions have remained challenging due to an incomplete understanding of underlying demographic processes (Fey et al., 2015). Jellyfish blooms are a prominent example for mass occurrences in which the population density of one life stage, the pelagic medusa, shows distinct peaks over broad temporal and regional scales (Lucas, Graham, & Widmer 2012). These demographic events may have lasting ecological, economic and social implications (cf. Lucas & Dawson, 2014). Jellyfish blooms can trigger trophic cascades such as collapse of pelagic food webs due to their severe predatory impact on a broad spectrum of zooplankton species and fish larvae (Richardson, Bakun, Hays, & Gibbons 2009; Riisgård, Andersen, & Hoffmann 2012; Schneider & Behrends, 1994, 1998). Further, jellyfish mass occurrences can interfere with human activities including tourism, fisheries, aquaculture and power production (Purcell, 2012; Purcell, Uye, & Lo 2007). Jellyfish populations, i.e. local aggregations of the medusa stage, seem to be increasing in numerous coastal ecosystems around the world (Brotz, Cheung, Kleisner, Pakhomov, & Pauly 2012). Available evidence suggests that jellyfish outbreaks which are defined as exceptional, sudden or ‘unnatural’ increases in biomass (Lucas & Dawson, 2014), may be favored by anthropogenic influences such as overfishing, eutrophication, habitat modification, species translocation and climate change (Purcell, 2012; Richardson, Bakun, Hays, & Gibbons 2009).

Bloom-forming jellyfish species are primarily found within the cnidarian taxon Scyphozoa and can feature complex life histories (Dawson & Hamner, 2009; Hamner & Dawson, 2009; Lucas & Dawson, 2014). Their metagenic life cycles typically include a pelagic, sexually reproducing medusa and a benthic, asexually reproducing polyp stage (Lucas, Graham, & Widmer 2012). Such life cycles are characterized by high stage-specific plasticity to changing environmental conditions (e.g. Gröndahl, 1989; Hamner & Jenssen, 1974; Liu, Lo, Purcell, & Chang 2009). Although the development of medusa populations is directly linked to survival rates of several precedent life stages, i.e. planula larvae, polyps and ephyrae (Lucas, 2001), information on the demographic rates of bloom-forming jellyfish has remained scarce (Xie, Fan, Wang, & Chen 2015). Quantifying demographic rates of all stages in the life cycle of species showing mass occurrence is thus fundamental to identifying the environmental variables causing changes in population dynamics and structure.

Here, we explore the demographic structure and environmental drivers underlying mass occurrence of the moon jellyfish *Aurelia aurita* (sensu stricto). We quantify demographic rates of all life stages of *A. aurita* (larvae, polyps, ephyrae and medusae), i.e. survival, stage-transitions and fecundity, based on experimental and field data from two temperate Danish water systems with contrasting prey availability. Using these demographic rates, we parameterize stage-structured population models to project the life cycle of jellyfish populations for low and high food levels (i.e. zooplankton prey biomasses). Since warm temperature has been associated with increased numbers of medusae for most temperate scyphozoan species (Purcell, 2005), we further predict the impact of a winter warming scenario on the population dynamics of *A. aurita* by perturbing demographic rates. Our results provide novel insights into the development of metagenic jellyfish populations under increased food levels and increased winter temperatures. Such information is essential to assess the boosting potential of environmental drivers on species showing mass occurrence and provides a milestone for identifying and managing demographic irregularities in marine ecosystems.

## Materials and Methods

### Study organism

The moon jellyfish *Aurelia* (Scyphozoa, Cnidaria) belongs to an ubiquitous, well-studied genus with at least 16 species, most of which are distributed in specific coastal and shelf sea regions around the world (Dawson, Gupta, & England 2005). The life history, i.e. individual stage-trajectory, of *Aurelia aurita*, sensu stricto, is characterized by a complex, metagenic life cycle which involves sexually reproducing medusae and asexually reproducing polyps (Figure 1a). Medusae of *A. aurita* are pelagic filter feeders which differentiate into distinct male and female individuals upon sexual maturation (Lucas 2001). After fertilization of oocytes, female medusae produce large numbers of lecithotrophic (yolk-feeding) planula larvae (Ishii & Takagi, 2003), which spend 12 h to 1 week in the water column (Lucas, 2001) before they settle on suitable hard substrate and metamorphose into polyps (Holst & Jarms, 2007; Keen, 1987). The inconspicuous benthic polyp stage of *A. aurita* can occur in densities between 60,000 and 400,000 ind. m^-2^ (Lucas, 2001) and reproduces asexually by several modes, including i) the production of well protected chitin-covered resting stages called podocysts (Arai, 2009), ii) budding of new polyps (Ishii & Watanabe, 2003), and iii) the release of multiple immature medusae, i.e. ephyrae, through transverse fission (strobilation). After asexual propagation, polyps remain in the polyp stage, while the development of pelagic ephyrae into sexually mature medusae closes the life cycle (Lucas, Graham, & Widmer 2012).

**Figure 1.**
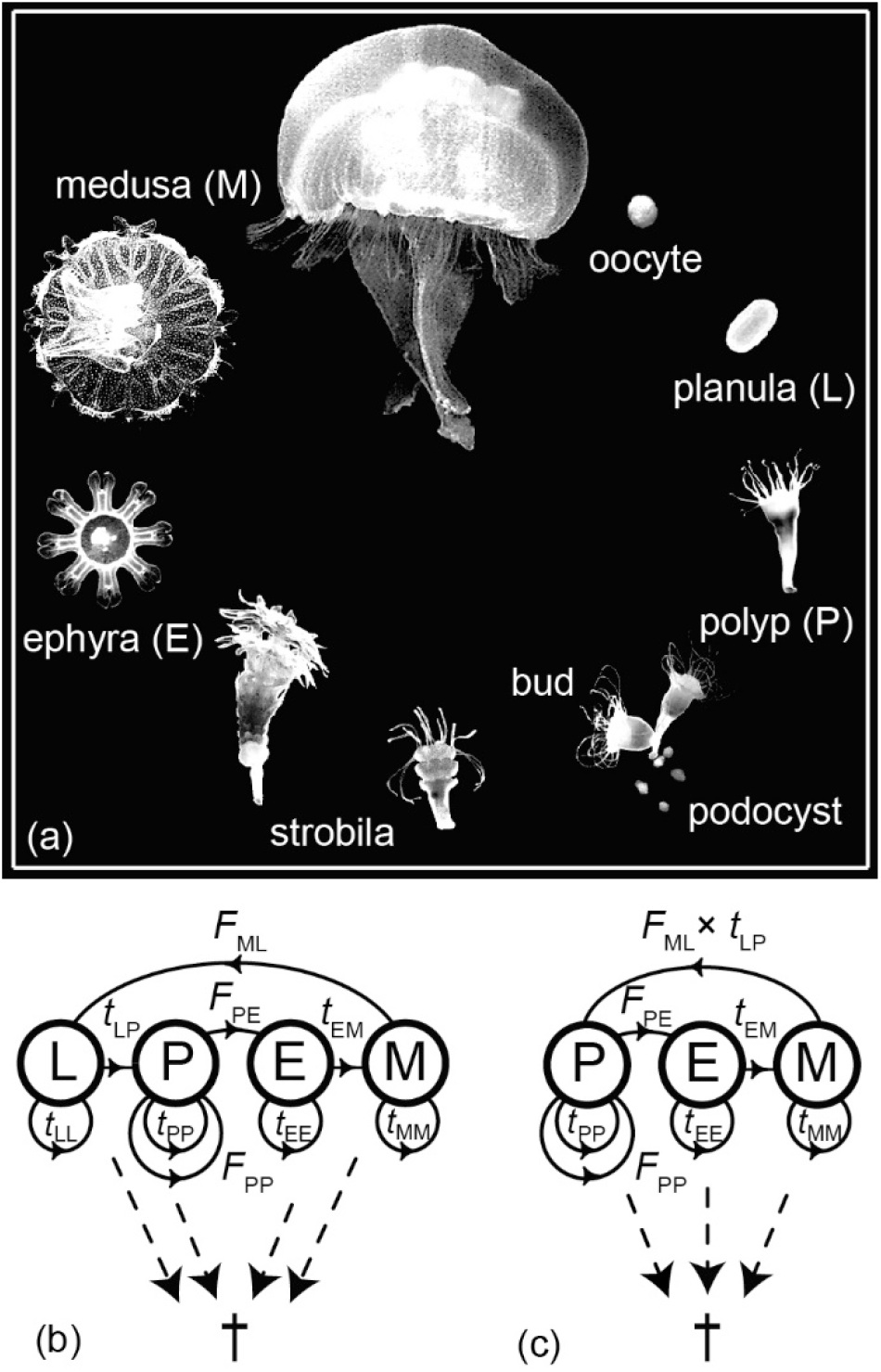
Life history of the moon jellyfish *A. aurita* (Scyphozoa, Cnidaria). (a) Metagenic life cycle with alternation of generations between the sexually reproducing medusa and the asexually reproducing polyp stage. (b) Life cycle graph for *A. aurita*, including transition probability *t*, fecundity *F* (solid arrows) and mortality † (broken arrows) of all life stages, i.e. planula larva (L), polyp (P), ephyra (E) and medusa (M). (c) Reduced life cycle graph for polyps, ephyrae and medusae considering fast (daily) transition from larva to polyp stage.

### Data collection

We cultivated planula larvae (L), polyps (P), ephyrae (E) and medusae (M) of *A. aurita* in the laboratories of the Marine Biological Research Centre, Kerteminde (Denmark) between June 2013 and November 2014 (Table 1). We set up separate experimental series for each life stage using filtered (38 µm) seawater at constant salinity (20 PSU) and with water temperature following seasonal variability of the local temperate climate zone (5-22 °C). We fed all life stages, except the planula larvae, with low (*L*) or high (*H*) amounts (corresponding to a ratio ∼1:10) of two-day old *Artemia salina* nauplii and regularly exchanged the seawater in our cultivation containers. We adjusted the fed biomass of *A. salina* (2.55 µg C ind.^-1^; Vanhaecke & Sorgeloos, 1980) to the size-specific energy demands of each life stage. In the following, we describe in more detail the origin and experimental setup for each investigated life stage.

**Table 1.** Experimental conditions for life stages of the jellyfish *Aurelia aurita* (L: planula larva, P: polyp, E: ephyra, M: medusa) under low (*L*) and high (*H*) food conditions. *l*_0_: initial number of individuals per treatment, *t*_0_, Δ*t*: start and duration of experiment, *c*: food concentration based on the carbon content of available prey organisms (*Artemia salina*).

| Food treatment | Life stage | $l_0$<br>(ind.) | $t_0$<br>(month) | $\Delta t$<br>(d) | $c$<br>($\mu\text{g C L}^{-1} \text{ d}^{-1}$) |
| --- | --- | --- | --- | --- | --- |
| *L* | L | 960 | Sept | 52 | 0 |
|  | P | 100 | Aug | 364 | 1 |
|  | E | 33 | Mar | 153 | 10 |
|  | M | 16 | Apr | 296 | - |
| *H* | L | 960 | Jun | 48 | 0 |
|  | P | 100 | Aug | 364 | 10 |
|  | E | 33 | Jan | 158 | 100 |
|  | M | 33 | Feb | 391 | >1000 |

We obtained planula larvae from female medusae that were collected in two contrasting Danish water systems – the fjord system Kerteminde Fjord/Kertinge Nor and the major water system Limfjorden. Zooplankton prey biomasses are in a range <1−100 µg C L^-1^ in Kerteminde Fjord/Kertinge Nor (Nielsen, Pedersen, & Riisgård 1997) and >10−1000 µg C L^-1^ in Limfjorden, respectively (Møller & Riisgård, 2007). Directly after the release of planulae, we transferred the larvae individually to multi-well seawater containers (200 µL). During daily counts, we determined the number of living larvae (*l*_LL_, ind. day^-1^) and the number of larvae that had completed metamorphosis into polyps (*l*_LP_, ind. day^-1^) until all planulae had metamorphosed into polyps or had died.

We cultivated polyps of *A. aurita* from planulae, released by female medusae from Kertinge Nor, which we initially added to small seawater containers (50 mL) with floating polystyrene substrate plates to facilitate settlement. After 24 h, we removed all but one newly settled polyp per substrate plate and replaced the seawater in the containers. We subsequently raised and cultivated the individual polyps under low or high food conditions (Table 1; cf. Kamiyama, 2011). To maintain the initial polyp cohorts, we instantly removed any buds of the “mother” polyps from substrate plates. During the whole experimental period of 12 months, no excystment of polyps from podocysts was observed. Each week we counted the number of live polyps (*l*_PP_, ind. week^-1^) as well as the number of buds (*N*_PP_, ind. week^-1^) and ephyrae per polyp (*N*_PE_, ind. week^-1^).

Ephyrae which were released by polyps under low and high food conditions were individually transferred to seawater flasks (250 mL) with constant water movement and air supply, and continuously reared under either low or high food conditions. Food levels for ephyrae corresponded to the minimum energy demands for maintenance of a ‘standard’ ephyra with 8 mm inter-rhopalia diameter (Båmstedt, Lane, & Martinussen 1999; Bengtson, Leger, & Sorgeloos 1991; Eckert, Randall, & Augustine 1988; Frandsen & Riisgård, 1997), or the 10-fold carbon load, respectively (Table 1). Each week we counted the number of surviving ephyrae (*l*_EE_, ind. week^-1^) and the number of ephyrae that had completed transition to medusa (*l*_EM_, ind. week^-1^); the latter we defined by the presence of fully developed intermediate lappets and the final closure of adradial clefts. We performed these counts until all ephyrae had either developed into medusae or had died.

We continuously tracked all newly transitioned medusae and additional sexually mature females collected in Kertinge Nor, in individual aquaria (1 L) with constant air supply. We kept the medusae under low or high food levels which resembled complete starvation or the 10-fold energy demands of a ‘standard’ 50 mm umbrella diameter medusa (Table 1; Olesen, Frandsen, & Riisgård 1994; Goldstein & Riisgård, 2016). We recorded the survival of medusae (*l*_MM_, ind. day^-1^) daily until all individuals in the low food treatment had died; at that time all medusae in the high food treatment were still alive. We estimated the daily release of planula larvae by female medusae during the reproductive period (*N*_ML_, ind. day^-1^) using the field observation-based relationship *N*_ML_ = 0.087 × 160.8 × e^0.029^*^d^* (Goldstein & Riisgård, 2016), where average umbrella diameters (*d*, mm) of 40 mm and 200 mm reflected low and high food conditions.

### Estimating stage-specific demographic rates

We used the experimental data to estimate survival rates, stage transition rates and fecundity of the stages L, P, E and M between age *x* and age *x*+1 for discrete time steps of one month. Individuals in each life stage *i* have certain monthly probabilities to survive in their current stage (*t_ii_*), to transition to another stage *j* (*t_ij_*) and to produce offspring of stage *j* (*F_ij_*). Each stage has its unique survival rate (*t*_LL_, *t*_PP_, *t*_EE_, *t*_MM_, month^-1^) and stage transitions are possible from larva to polyp (*t*_LP_, month^-1^) or from ephyra to medusa (*t*_EM_, month^-1^). Fecundity includes asexual reproduction of polyps by budding (*F*_PP_, ind. ind.^-1^ month^-1^) or release of ephyrae (*F*_PE_, ind. ind.^-1^ month^-1^), and sexual reproduction of medusae by the release of planula larvae (*F*_ML_, ind. ind.^-1^ month^-1^; Figure 1b). Details on estimating monthly stage-specific demographic rates of *A. aurita* are available from the Supporting Information (Appendix S1 & S2).

We first estimated monthly transition probabilities and fecundities for each life-stage *i* under low and high food conditions (Appendix S1) based on count ratios from our empirical data with either daily (L, M) or weekly resolution (P, E). For converting the daily release of planula larvae into monthly fecundity of medusae, we assumed a sex ratio of 1:1. Since transition probabilities and fecundities between subsequent time steps depend on survivorship within each monthly time step, we assumed that individuals that died during a given month, had died halfway during this month, i.e. they survived throughout half the month on average (Appendix S2). This assumption is needed as, for instance, a medusa might still release larvae before its death within a given month. Further, some ephyrae completed transition to the medusa stage within less than a month after they were released by a polyp. We therefore corrected the monthly release of ephyrae by polyps not only for daily survivorship of ephyrae, but also for the transition probability to the medusa stage within a given month.

### Stage-structured matrix population models

Our matrix population models describe the transitions of each life stage taking place during each projection interval, as defined by monthly stage-specific demographic rates for each of the 12 month of a year. Age is determined by month, and the final stages are P, E and M. Since larval transition to the polyp stage was restricted to days or hours (*t*_LP_ = 0 for incompetent larvae with *t*_LL_ >0), we linked the monthly transition from larva to polyp stage to fecundity of medusae (*F*_ML_×*t*_LP_) to build the basic structure of an irreducible monthly stage-structured matrix model (Figure 1c). Using our empirical datasets on all demographic rates of *A. aurita* under low and high food conditions, respectively, we constructed monthly stage-structured submatrices of the following structure:

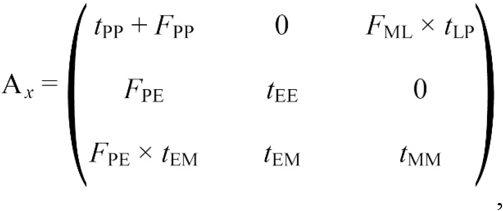

where *x* denotes the age in months. Based on 12 monthly submatrices A*_x_* (3 × 3) which reflect seasonal life cycle dynamics of temperate *A. aurita* populations (Appendix S3: Figures S1 & S2), we constructed yearly population matrices A (36 × 36; time step = 1 month) for monthly projections over a 10-year interval (120 iterations):

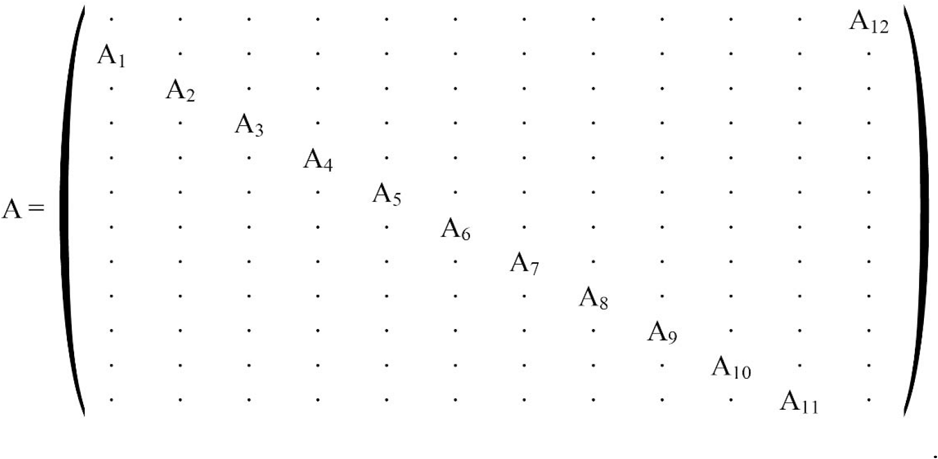

In some months, we observed no survivorship of ephyrae or medusae, which would lead to reducible matrices (Caswell, 2001); in these cases, we included a small transition probability *c* = 1 × 10^-10^ as placeholder (Appendix S3: Figures S1 & S2). Our results were robust to this assumption.

### Adjusting matrix parameters to environmental conditions

We constructed yearly projection matrix models for the jellyfish population under low and high food conditions. Since our transition probabilities from larva to polyp were unrealistically high due to the sheltered laboratory environment (as an important microzooplankton component, planula larvae are subject to a high mortality risk under natural conditions), we calibrated the larva to polyp transition to a level (*t*_LP_ = 0.001 month^-1^) that resulted in a stationary population (i.e. characterized by constant population size over time) under low food conditions. We used the same *t*_LP_ as for the calibrated low food condition to scale the projection matrix model for high food conditions, leaving us with a rapidly growing population.

We parameterized additional projection matrix models to explore the potential effects of a winter warming scenario under low and high food conditions. A simulated increase in present water temperatures from 5 to 10 °C is predicted to benefit the production of *A. aurita* ephyrae in terms of extended strobilation periods and enhanced ephyra release per polyp (Holst, 2012). Our winter warming scenario included 1.8-fold increased ephyra production (i.e. *F*_PE_×1.8) during strobilation periods that we extended by one month, i.e. mean *F*_PE_ of proceeding and subsequent month. We additionally adjusted corresponding probabilities for ephyra survival and transition from ephyra to medusa to *t*_EE_ and *t*_EM_ of subsequent month, respectively (Appendix S3: Figures S1 & S2). With these modified matrices, we investigated the demographic effects of a predicted winter warming trend on *A. aurita* populations under low and high food conditions.

### Computing demographic parameters and population projection

We analyzed our yearly population projection matrices using R version 3.1.3 (R Core Team, 2015) and compared the demographic dynamics of *A. aurita* under low and high food levels, and in combination with a winter warming trend. We used R packages ‘popbio’ (Stubben & Milligan, 2007) and ‘popdemo’ (Stott, Hodgson, & Townley 2012) to compute the following population-level metrics: population growth rate λ (dominant eigenvalue), stable stage distribution (right eigenvector **w** corresponding to the dominant eigenvalue; i.e. proportions of individuals in each stage remain constant on the long-term), reproductive value *R* (left eigenvector **v** corresponding to the dominant eigenvalue; i.e. expected lifetime reproduction of an individual in stage *i*), net reproductive rate *R*_0_ (i.e. average number of offspring by which an individual will be replaced during its lifetime), generation time *T* (i.e. average age at which an individual has produced the average number of offspring, *R*_0_), life expectancy *e_0_*(i.e. mean time to death), and sensitivity of λ to perturbations of the matrix elements (only elements >0 are considered). We further performed a fixed life table response experiment (LTRE) analysis (Caswell, 1996; Tuljapurkar & Caswell, 1997) for pairwise comparison of the sensitivity of λ to changes in stage-specific transitions under high versus low food conditions.

## Results

### Demographic dynamics of jellyfish under low and high food conditions

The metagenic life cycle of *A. aurita* (Figure 1a) was reflected by the cyclic (imprimitive) structure of our matrix model, expressing annual peak and pit dynamics in projected population density (Figure 2a). Relative to the calibrated stationary stable population with λ = 1 month^-1^ under low food conditions, the scaled population growth rate for 10-fold increased food availability was λ = 1.63 month^-1^, corresponding to an approximated increase in population size by a factor (λ^12^ =) 346 year^-1^ (Table 2). Such differences in growth rate due to changes in food availability provide high leverage for population booms and busts. Similar to low food levels, the projection matrix under high food availability showed cyclic annual structure with typical peak and pit dynamics in projected population density, but life cycle amplitudes were much increased (Figure 2b).

**Figure 2.**
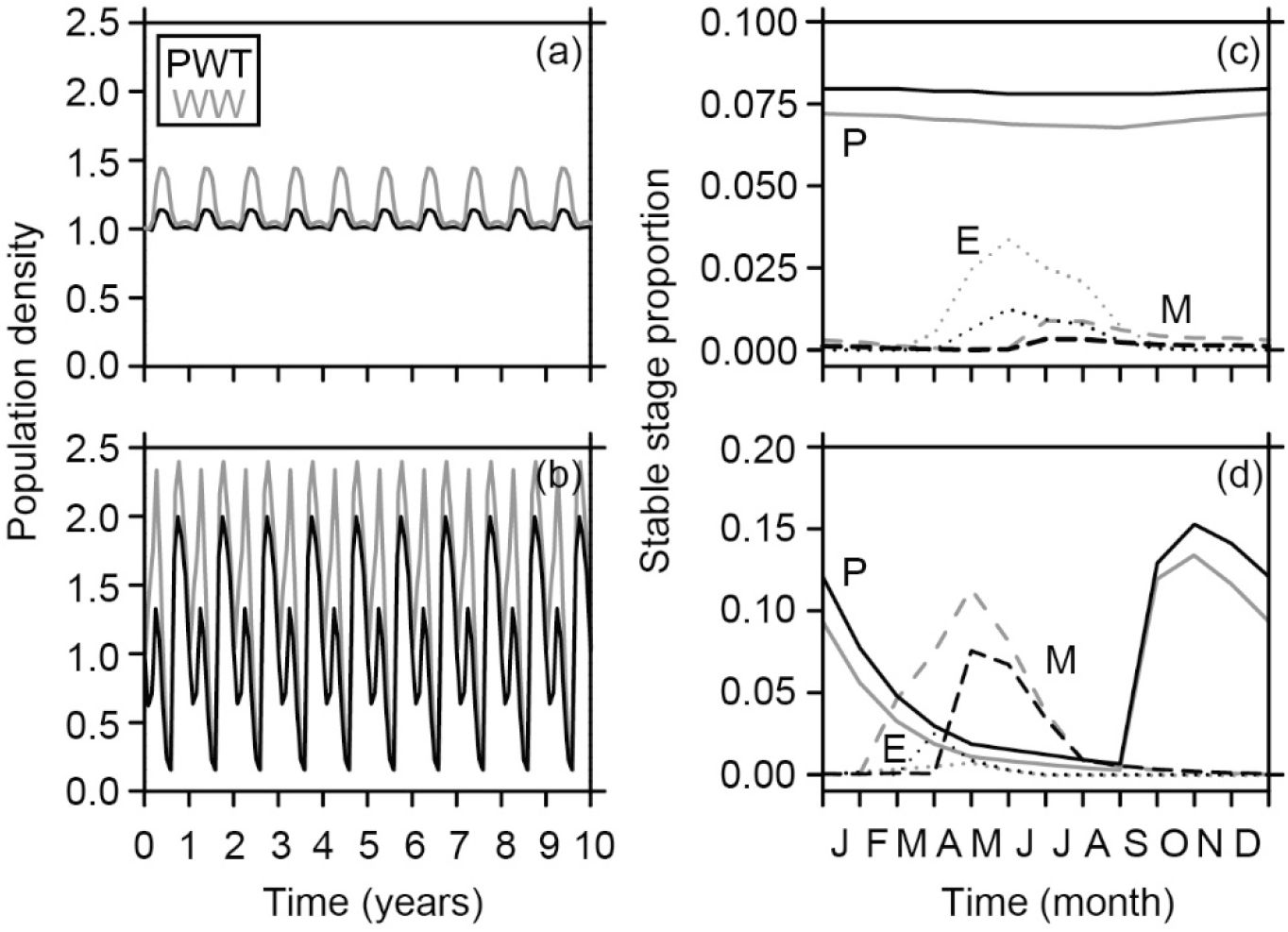
Population matrix model for *A. aurita* under low (upper panels) and high (lower panels) food conditions (∼1:10) at present water temperatures (PWT) and in a winter warming scenario (WW). (a−b) Matrix projection over a 10-year period, standardized to the long-term effects of population growth rates λ = 1 month^-1^ (low food availability) and λ = 1.63 month^-1^ (high food availability), respectively. (c−d) Stable stage distribution (i.e. relative, constant long-term proportions) of polyps (P; solid lines), ephyrae (E; dotted lines) and medusae (M; broken lines).

**Table 2.** Demographic parameters of yearly stage-structured projection matrices (P: polyp, E: ephyra, M: medusa) for a local *Aurelia aurita* population under low (*L*) and high food conditions (*H*) at present water temperatures (PWT) and for a predicted winter warming trend (WW).

| Demographic parameter | *L* |  | *H* |  |
| --- | --- | --- | --- | --- |
|  | PWT | WW | PWT | WW |
| Scaled population growth rate $\lambda$ (month<sup>-1</sup>) | 1.00 | 1.01 | 1.63 | 1.73 |
| Damping ratio $\rho$ (months) | 58.6 | 59.2 | 2.8 | 3.2 |
| Stable stage proportion (P, E, M) | 0.94, 0.04, 0.02 | 0.84, 0.12, 0.04 | 0.76, 0.04, 0.20 | 0.60, 0.02, 0.38 |
| Scaled reproductive value $R$ (ind.) (P, E, M) | 1, 0.03, 0.42 | 1, 8.53, $3 \times 10^{-7}$ | 1, 0.27, 9.13 | 1, 0.18, 7.43 |
| Net reproductive rate $R_0$ (ind.) | 1.0 | 1.8 | 57.0 | 80.9 |
| Generation time $T$ (months) | 200.0 | 111.8 | 8.3 | 8.0 |
| Life expectancy $e_0$ (months) (P, E, M) | 394.5, 1.7, 3.1 | 394.5, 2.1, 3.1 | 86.7, 2.9, 8.0 | 86.7, 3.5, 8.0 |

Under low food conditions, the overall population density (taking into account all life stages) increased to a peak during spring and summer and thereafter decreased to low densities during autumn and winter. Within-year dynamics under high food levels differed significantly compared to low food conditions, showing two distinct maxima in population density, one during spring and one during autumn, each followed by a subsequent minimum in summer and winter (Figures 2a−b). The convergence time to stable stage (i.e. the damping ratio ρ) of our modelled *A. aurita* population was ∼5 years under low food conditions and reduced by a factor 21, for the high food population (Table 2).

The stable stage distribution under low food availability was vastly dominated by polyps (0.94) that survived throughout the year. Ephyrae and medusae accounted for comparatively small fractions of the population (0.04 and 0.02, respectively) and showed typical seasonal patterns. Ephyrae were released during April/May and survived into September. Only few ephyrae transitioned to the medusa stage under low food conditions, so that medusae were present from June to December and contributed only marginally to restocking polyp densities by the release of planula larvae during autumn and winter (Figure 2c). Under high food supply, less polyps (0.76) dominated the overall *A. aurita* population and the stable stage distribution was characterized by a 10-fold increased medusa fraction (0.2) compared to low food conditions (Table 2). Compared to a year-round standing stock of polyps under low food conditions, the stable stage proportion of polyps under high food levels showed a peak in autumn after being restocked by the release of planula larvae, and subsequently decreased throughout the year. Ephyrae were released in March/April, and the medusa fraction dominated the *A. aurita* population from April to July under high food levels (Figure 2d).

The reproductive value was low for ephyrae (0.03) under low food conditions, and sexual reproduction of medusae contributed only about half as much (0.42) to future generations as the asexually reproducing polyp stage (1; note scaling to 1 for polyps in January). Under high food conditions, the reproductive value of ephyrae and medusae increased by a factor 8.1 and 21.6, respectively (Table 2). A net reproductive rate of 1 indicated that each polyp is replaced by 1 new polyp at the end of its life under low food conditions which resulted in long generation times of ∼16.7 years. High food levels increased the net reproductive rate to 57 polyps replacing each polyp at the end of its life and boosted the average time between two consecutive generations to 8 months. Life expectancy of polyps, estimated as the mean time to death, was 33 years under low food conditions, while ephyrae and medusae revealed much shorter life expectancies of 2 and 3 months, respectively. In consensus with a more pronounced impact of the medusa stage on population structure under high food levels, life expectancy of polyps decreased to 7 years, while the length of ephyra and especially medusa lives extended to 3 and 8 months, respectively (Table 2).

### Sensitivity of demographic rates to increased food availability

Sensitivity analysis generally highlighted a strong influence of survival rates and less pronounced effects of fecundity on the growth of jellyfish populations. Under low food conditions, population growth was most sensitive to changes in polyp survival (*t*_PP_+*F*_PP_), expressed by constant sensitivity 0.08 (relative maximum value in the sensitivity matrix) throughout the months of the year (Figure 3a). Comparatively, λ showed a much lower sensitivity of 0.02 to changes in ephyra production (*F*_PE_) from March to May. The overall sensitivity of λ to perturbations in the projection matrix elements increased under high compared to low food levels. Under high food conditions, changes in polyp survival (*t*_PP_+*F*_PP_) affected λ particularly from October to April, with sensitivities of 0.06 to 0.13. In contrast to low food levels, however, λ was most sensitive to changes in medusa survival (*t*_MM_) under high food levels. Sensitivity to *t*_MM_ remained high throughout the year, ranging from 0.07 to 0.16 from April to October, and reaching a maximum of 0.32 from June to July (Figure 3b). We further detected sensitivities of λ in the range 0.02 to 0.03 for perturbations in ephyra production (*F*_PE_) from March to May under high food conditions. Differences in the sensitivity of λ between food treatments were also evident from the results of our life table response experiment (LTRE) analysis. High food availability led to 7.7 to 12.6 % increased sensitivity for the transition probabilities from polyp to medusa stage (i.e. *F*_PE_+*t*_EM_) from March to May. Further, increased food conditions enhanced the sensitivity of λ for medusa survival (i.e. *t*_MM_) during April by 7.7 % and for polyp production via sexual reproduction of medusae (i.e. *F*_ML_+*t*_LP_) from August to December by 5.0 to 6.8 % (Figure 3c).

**Figure 3.**
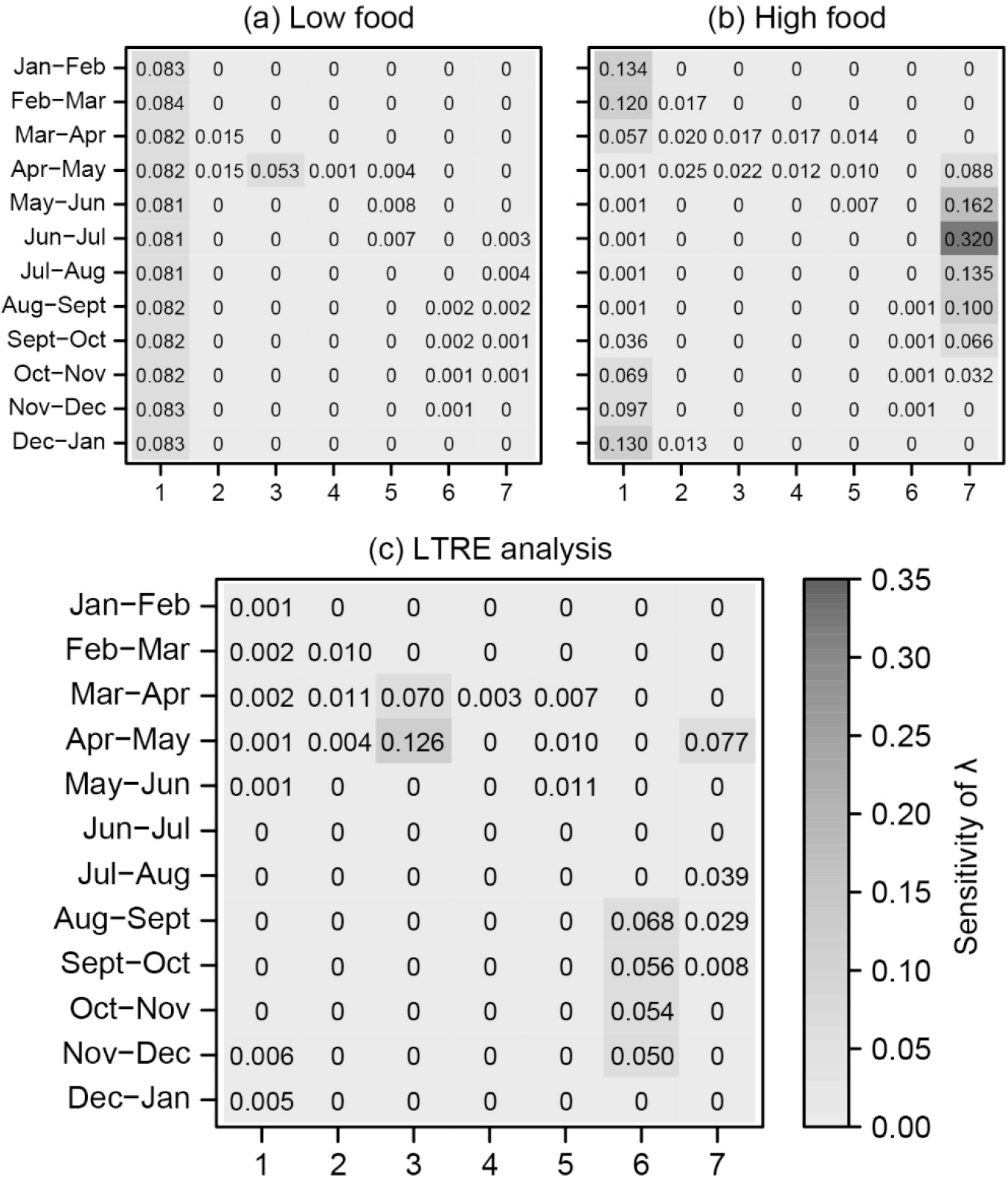
Sensitivity of the population growth rate λ of *A. aurita* to changes in monthly stage-specific transitions (1: *t*_PP_+*F*_PP_, 2: *F*_PE_, 3: *F*_PE_×*t*_EM_, 4: *t*_EE_, 5: *t*_EM_, 6: *F*_ML_×*t*_LP_, 7: *t*_MM_) under (a) low and (b) high food conditions. (c) Life table response experiment (LTRE) analysis for sensitivity of λ to perturbations in the population projection matrix elements when comparing high to low food conditions (∼10:1).

### Demographic dynamics of jellyfish in a winter warming scenario

For our winter warming scenario with extended strobilation periods and increased ephyra release, population growth rates of *A. aurita* increased to 1.01 and 1.73 month^-1^ under low and high food conditions, corresponding to 1.1- and 729-fold yearly increases in population size (i.e., relative increases by 6.2 % and 110.9 %), respectively. Winter warming led to stronger variability in inter-annual population structure in both food treatments, expressed by more distinct peaks and pits in population density (Figures 2a−b). Under low food levels, the stable population structure of *A. aurita* consisted of 2.7 % more ephyrae, 7.9 % more medusae, and 10.6 % less polyps per year as a consequence of winter warming (Figure 2c). Under high food levels, increased winter temperatures resulted in a stable stage distribution with 17.4 % more medusae, while the proportions of ephyrae and polyps decreased by 1.7 % and 15.7 %, respectively (Table 2). An earlier onset of strobilation periods in the scenario and fast transition of ephyrae caused significant proportions of medusae to be already present from February (Figure 2d).

Winter warming caused substantial changes in reproductive rates and generation times of the projected jellyfish populations; these changes were particularly pronounced under low food conditions. The reproductive value of ephyrae in the low food population was 257-fold increased, while the relative contribution to the number of offspring by medusae decreased to near zero. Winter warming in combination with high food levels decreased reproductive values of both ephyrae and medusae by a factor 1.5 and 1.2, respectively (Table 2). Increased winter temperatures raised the net reproductive rate by 75.1 % under low food conditions and by 42.0 % under high food conditions. The winter warming scenario shortened the generation time by 7.4 years under low food conditions, while generation times were shortened by only 9 days under high food conditions. Winter warming increased the life expectancy of ephyrae by 12 and 18 days, respectively, whereas life expectancy of polyps and medusae remained unchanged under both low and high food levels (Table 2).

## Discussion

We reveal the demographic dynamics and life history shifts underlying jellyfish mass occurrence and show that variable intensity and frequency is subject to specific ecological triggers. Our matrix projections quantify how jellyfish blooms are enhanced by naturally occurring variations in the food regime. We suggest the observed temporal shift from polyp- to medusa-dominated populations in response to increased food availability as a key mechanism causing jellyfish outbreaks, which can be boosted by the combined effects with a predicted winter warming. Our findings support previous studies which have pointed out metagenic life cycles as an ultimate driver of boom and bust population dynamics (Brotz, Cheung, Kleisner, Pakhomov, & Pauly 2012; Hamner & Dawson, 2009; Lucas & Dawson, 2014, their Figure 2.4b; Pitt, Welsh, & Condon 2009). Considering jellyfish outbreaks as highly sensitive indicators for ocean degradation (Schrope, 2012), the present insights may provide fundamental knowledge regarding the health of marine ecosystems worldwide.

### Food availability as a key driver of jellyfish blooms

Our study provides insight into probable consequences of habitat eutrophication in context with jellyfish blooms, as investigated food levels mirror eutrophic to hypereutrophic conditions prevailing in several Danish coastal regions (Riisgård, Jensen, & Rask 2008, their Table 16.1). Resulting demographic dynamics suggest that food availability is a major constraint for the intensity and frequency of jellyfish blooms, confirming previous observations of food availability as a key variable in limiting the development of ephyrae (Fu, Shibata, Makabe, Ikeda, & Uye 2014) and the survivorship and fecundity of medusae (Goldstein & Riisgård, 2016). Yet, to our knowledge, we present the first comprehensive quantitative model of how increased food availability can trigger demographic changes in a scyphozoan jellyfish population. Such changes include enhanced boom and bust dynamics and shorter generation times. We show that complex life cycles can facilitate jellyfish blooms and emphasize the key role of benthic polyps in ensuring long-term survival (Boero et al., 2008; Lucas, Graham, & Widmer 2012), particularly when food is scarce. Life-history trade-offs such as extended polyp life-spans along with reduced rates of asexual reproduction under low food levels (Stearns, 1976) may explain high plasticity of jellyfish populations to changing environmental conditions. In the moon jellyfish, increased food levels are associated with boosted population growth rates and dramatic life history shifts, as expressed by longer-lived medusae with enhanced release of planula larvae, increased ephyra production, as well as faster and more successful development of ephyrae into medusae.

The availability of food resources further drives a population density-dependent mechanism controlling individual size (Lucas, 2001) and thus, biomass, which plays a major role for the ecological impact of *A. aurita* (Riisgård, Barth-Jensen, & Madsen 2010; Schneider & Behrends, 1994). Size-specific traits should hence be taken into account for future population models (cf. Bordehore et al., 2015) to quantify and evaluate the development of jellyfish blooms from a food web perspective (cf. Schnedler-Meyer, Kiørbøe, & Mariani 2018). Although the temporal and spatial patterns of abundance in each life stage may vary according to jellyfish species (Lucas, Graham, & Widmer 2012; Scorrano, Aglieri, Boero, Dawson, & Piraino 2016) and even populations of the same species (Dawson et al., 2015), our observations on the moon jellyfish are likely describing general trends. Field evidence of a strong positive correlation between medusa density and prey abundance for two distinct *A. aurita* populations (Dawson et al., 2015, their Figure 4d) implies that observed population outbreaks in response to increased food supply are common across the *Aurelia* species complex and other metagenic jellyfish species.

### Winter warming and jellyfish blooms

Our projected winter warming scenario indicates pronounced effects of an increase in water temperatures from 5 to 10 °C on the development of temperate *A. aurita* populations in combination with regionally observed increases in food level. Based on recent climate analyses, rises in global ocean surface temperature by 1 to 2 °C can be expected within the next 100 years, with a particularly strong winter warming trend in semi-enclosed ecosystems such as the North and Baltic Seas (Belkin, 2009; Mackenzie & Schiedeck, 2007). As reflected by doubled yearly population growth rates under high food conditions in response to the simulated increase in winter temperatures, our findings support that increased water temperature can boost the future booms and busts of jellyfish blooms (cf. Richardson, Bakun, Hays, & Gibbons 2009; Xie, Fan, Wang, & Chen 2015). According to our population matrix model, underlying seasonal changes in population composition and structure due to winter warming include a general trend towards more medusae and less polyps under increased food conditions. In regions with limited food availability, winter warming can further increase the net reproductive rate of temperate jellyfish populations. Due to enhanced strobilation periods and increased numbers of released ephyrae per polyp (Holst, 2012; Purcell, 2005) as simulated in our matrix model projections, warmer winter temperatures may shorten the generation time under low food conditions drastically.

Climate warming has previously been pointed out to benefit the population size of temperate jellyfish species, but not tropical or boreal species (Purcell et al., 2012). Further, a cold trigger is necessary to stimulate strobilation in many scyphozoans (Holst, 2012; Lotan, Fine, & Ben-Hillel 1994), while heatwave events can negatively affect the population dynamics of jellyfish (Chi, Mueller-Navarra, Hylander, Sommer, & Javidpour 2018). Taking into account these physiological constraints, we suggest winter warming may promote the blooming potential of certain temperate jellyfish species but seems to be a less powerful driver when compared to increased food availability.

### Jellyfish life histories in a changing environment

Beyond revealing the demographic key variables that control population growth of *A. aurita* in response to habitat eutrophication and climate warming, our matrix model projections provide general implications for the development of jellyfish blooms with regard to environmental change. In the following, we address how other potential drivers of jellyfish blooms which have not been quantified in the present study can overrule important ecological parameters that constrain the demography of metagenic jellyfish. Such natural limitations include intra- and inter-specific competition, predation, substrate availability, hydrodynamic currents and the effects of salinity, hypoxia, pH, pollution, light and sedimentation (Arai, 2005; Johnson, Perry, & Burke 2001; Lucas, Graham, & Widmer 2012).

Our findings emphasize the crucial role of predation by benthic fauna, spatial competition and availability of settlement substrate for transition from the planktonic planula to the benthic polyp stage under natural conditions (Lucas, 2001; Lucas, Graham, & Widmer 2012). We estimated a transition probability of 0.001 month^-1^ from larva to polyp, which is in agreement with previous estimates for *A. aurita* (Xie, Fan, Wang, & Chen 2015), when scaling to a stable *A. aurita* population under low food levels (λ = 1 month^-1^). The local *A. aurita* jellyfish population in the semi-enclosed fjord system Kerteminde Fjord/Kertinge Nor, where zooplankton biomasses are comparable to our low food conditions, has remained unchanged over the last 25 years (Goldstein & Riisgård, 2016; Olesen, Frandsen, & Riisgård 1994; Riisgård, Barth-Jensen, & Madsen 2010) and may, similar to our projected *A. aurita* population under low food conditions, mainly depend on environmental parameters that determine larval transition to the benthic polyp stage.

The rapid growth potential we show for our calibrated jellyfish population under constantly high food (optimum) conditions however stands in contrast to rather low densities of *A. aurita* medusae observed in open coastal waters (Goldstein & Riisgård, 2016; Möller, 1980) or in Limfjorden (Hansson, Moeslund, Kiørboe, & Riisgård 2005; Møller & Riisgård, 2007), which implies additional factors regulating the demographic dynamics of jellyfish in less protected ecosystems. The results of our life table response experiment analysis suggest that besides successful metamorphosis of planula larvae, transition from ephyra to medusa and survival of medusae are the most critical parameters for the development of *A. aurita* populations in exposed coastal regions. According to our matrix projection for high food levels, reduced medusa survival (*t*_MM_ = 0.1 month^-1^), as a possible consequence of natural predation by other cnidarians, ctenophores or fish (Arai, 2005), or alternatively, a lowered transition probability from ephyra to medusa stage (*t*_EM_ = 0.0001 month^-1^), would remarkably decelerate the population growth rate (from λ =1.63 month^-1^ to 1.10 month^-1^ and 1.09 month^-1^, respectively).

Since jellyfish constitute major parts of the diet of some fish species (Arai, 2005; Hamilton, 2016) and can in turn be important competitors for zooplankton prey with forage fish (Purcell, 2012), these findings indicate dramatic effects of large-scale alterations in ecosystem dynamics by overfishing on the demography of the pelagic medusa and ephyra stages. Lack of competition and predation by fish may further create protected niches for nonindigenous jellyfish species that are inter-regionally transported via ballast water exchange (Richardson, Bakun, Hays, & Gibbons 2009). Man-made shoreline modifications contribute decisively to the availability of hard substrate for benthic polyps (Purcell, 2012) and are, according to our findings, likely to enhance the impact of changing food web structures and climate conditions on jellyfish blooms. For these reasons, we expect that future jellyfish populations will further exploit their growth potential in the context of environmental change. Removal of jellyfish and reduction of their prey availability have previously been considered as possible management strategies (Bordehore et al., 2015) which, however, could have unpredictable consequences for the sensitive balance of marine food webs. A decisive step forward will be to further explore jellyfish life histories in stochastic environments, taking into account specific traits that facilitate survival of metagenic species under unfavorable conditions, such as production of resting stages (podocysts and planulocysts), regeneration and reverse development (Piraino, De Vito, Schmich, Bouillon, & Boero 2004).

## Conclusions

Our study highlights that food availability can drive and specifically shape the occurrence of jellyfish blooms. We suggest that for most temperate scyphozoans predicted changes in water temperature may cause less dramatic increases in jellyfish density than already established food concentrations in several eutrophic regions around the world. The role of natural competitors and predators in controlling jellyfish populations needs further exploration, but our findings might serve as a promising starting point. We conclude that especially the combination of key environmental drivers, such as habitat eutrophication and climate change, may severely promote jellyfish blooms. Our findings emphasize the fundamental importance of studying jellyfish life histories using quantitative approaches. Such approaches, which take into account species-, life stage- and age-specific variation in demographic rates, are highly valuable tools for informing future management of mass occurrences in marine ecosystems.

## Authors’ contributions

US and JG conceived the ideas and designed the methodology, JG collected the data and led the writing of the manuscript; JG and US analyzed the data. Both authors contributed critically to the drafts and gave final approval for publication.

## Acknowledgements

We thank Kim Lundgreen for technical assistance and Daniel A. Levitis, Hans Ulrik Riisgård, Hal Caswell, Iain Stott, Cesar Bordehore, Per Andersen, Owen Jones, Matthew Spencer and Cathy H. Lucas for helpful comments and discussions. We are grateful to the reviewers for providing valuable feedback on the manuscript. This study was financially supported by the Max-Planck Society (Max-Planck Odense Center on the Biodemography of Aging, Denmark). The authors declare no conflict of interest.

## Data Accessibility

A preprint version of this article is available from bioRxiv https://doi.org/10.1101/102814 (Goldstein & Steiner, 2019). Data will be made publicly accessible on the Dryad Digital Repository at the time of publication.

## Supporting Information

Additional Supporting Information is available online in the supporting information tab for this article.

