## Supporting Information for "Ecological drivers of jellyfish blooms – the complex life history of a ‘well-known’ medusa (*Aurelia aurita*)"

J. Goldstein* [a, b], U.K. Steiner [b, c]

[a]Marine Biological Research Centre, University of Southern Denmark, Hindsholmvej 11, 5300 Kerteminde, Denmark

[b]Department of Biology, University of Southern Denmark, Campusvej 55, 5230 Odense, Denmark

[c]Center on Population Dynamics, University of Southern Denmark, Campusvej 55, 5230 Odense, Denmark

*Journal of Animal Ecology*

---

### S1 Stage-specific transition probabilities

For each stage *i*, we calculated the stage-specific monthly survival probability in stage *i* from age *x* to age *x*+1 (${}_{x+1}t_{ii,x}$, month$^{-1}$) using the equation:

$$ {}_{x+1}t_{ii,x} = \frac{l_{ii,x+1}}{l_{i,x}} \quad \text{(S1)}, $$

where $l_{ii,x+1}$ (ind.) is the number of individuals surviving in stage *i* from age *x* to age *x*+1 and $l_{i,x}$ (ind.) the number of live individuals in stage *i* at age *x*. The monthly probability of death for each stage *i* between ages *x* and *x*+1 (${}_{x+1}q_{i,x}$, month$^{-1}$) is defined as

$$_{x+1}q_{i,x} = \frac{d_{i,x+1}}{l_{i,x}} \quad \text{(S2)},$$

where $d_{i,x+1}$ (ind.) describes the number of individuals dying in stage $i$ from age $x$ to age $x$+1. Monthly transition probability from stage $i$ to stage $j$ in the age interval $x$ to $x$+1 ($_{x+1}t_{ij,x}$, month$^{-1}$) was determined as follows:

$$_{x+1}t_{ij,x} = \frac{l_{ij,x+1}}{l_{i,x}} \quad \text{(S3)},$$

where $l_{ij,x+1}$ is the number of individuals that completed transition from stage $i$ to stage $j$ between age $x$ and age $x$+1. From the total number of offspring produced by each stage $i$ asexually or sexually ($N_{ii/ij}$, ind. month$^{-1}$), we calculated monthly fecundity ($F_{ii/ij}$, ind. ind.$^{-1}$ month$^{-1}$) in the age interval $x$ to $x$+1 as

$$_{x+1}F_{ii/ij,x} = \frac{\sum N_{ii/ij,x+1}}{l_{i,x}} \quad \text{(S4)}.$$

#### S2 Survival-dependent correction of stage-specific rates

Transition probabilities and fecundities take into account survivorship during each monthly time step based on the assumption that individuals that died during a given month, have lived during half the month on average. We therefore corrected the monthly transition rate from larva to polyp stage ($t_{LP}$) for polyp survival as follows:

$$t_{LP\_cor} = t_{LP} \times \left(1 - 0.5 \times q_P\right) \quad \text{(S5)},$$

with $q_P$ being the polyp mortality, and subscripts referring to the different life stages, i.e. larvae (L), polyp (P), ephyrae (E) and medusae (M). We also corrected monthly transition from ephyra to medusa stage ($t_{EM}$) by considering medusa survival:

$$t_{EM\_cor} = t_{EM} \times \left(1 - 0.5 \times q_M\right) \quad \text{(S6)}.$$

In a similar way we corrected the production of new polyps in the polyp stage ($F_{PP}$) for polyp survival during each month:

$$F_{PP\_cor} = F_{PP} \times \left(1 - 0.5 \times q_P\right) \quad \text{(S7)}.$$

As some ephyrae completed transition to medusa stage within less than a month after release by a polyp, the monthly release of ephyrae by polyps ($F_{PE}$) was corrected for survivorship of ephyrae and for the transition probability of survivors to the medusa stage as follows:

$$F_{\mathrm{PE\_cor}} = F_{\mathrm{PE}} - F_{\mathrm{PE}} \times t_{\mathrm{EM}} \times \left(1 - 0.5 \times q_{\mathrm{E}}\right) \qquad \text{(S8).}$$

We further corrected the fecundity of medusae, i.e. the production of planula larvae ($F_{\mathrm{ML}}$) for survivorship of larvae during each month using the equation:

$$F_{\mathrm{ML\_cor}} = F_{\mathrm{ML}} \times \left(1 - 0.5 \times q_{\mathrm{L}}\right) \qquad \text{(S9).}$$

### S3 Matrix construction

$$A_{L1} = \begin{bmatrix} 1 & 0 & 0 \\ 0 & c & 0 \\ 0 & 0 & 0.500 \end{bmatrix} \quad A_{L5} = \begin{bmatrix} 1 & 0 & 0 \\ 0 & 0.750 & 0 \\ 0 & 0.250 & 1 \end{bmatrix} \quad A_{L9} = \begin{bmatrix} 1 & 0 & 333 \\ 0 & c & 0 \\ 0 & 0 & 0.857 \end{bmatrix}$$

$$A_{L2} = \begin{bmatrix} 0.990 & 0 & 0 \\ 0^{0.074} & c & 0 \\ 0 & 0 & 0.500 \end{bmatrix} \quad A_{L6} = \begin{bmatrix} 1 & 0 & 0 \\ 0 & 0.833 & 0 \\ 0 & 0.039 & 0.875 \end{bmatrix} \quad A_{L10} = \begin{bmatrix} 1 & 0 & 344 \\ 0 & c & 0 \\ 0 & 0 & 1 \end{bmatrix}$$

$$A_{L3} = \begin{bmatrix} 1 & 0 & 0 \\ 0.082^{0.281} & c^{0.970} & 0 \\ 0^{0.005} & 0^{0.015} & c \end{bmatrix} \quad A_{L7} = \begin{bmatrix} 1 & 0 & 0 \\ 0 & 0.350 & 0 \\ 0 & 0 & 0.714 \end{bmatrix} \quad A_{L11} = \begin{bmatrix} 1 & 0 & 333 \\ 0 & c & 0 \\ 0 & 0 & 0.833 \end{bmatrix}$$

$$A_{L4} = \begin{bmatrix} 0.990 & 0 & 0 \\ 0.078^{0.141} & 0.970 & 0 \\ 0.001^{0.002} & 0.015 & c \end{bmatrix} \quad A_{L8} = \begin{bmatrix} 0.990 & 0 & 342 \\ 0 & 0.143 & 0 \\ 0 & 0 & 0.700 \end{bmatrix} \quad A_{L12} = \begin{bmatrix} 1 & 0 & 0 \\ 0 & c & 0 \\ 0 & 0 & 0.800 \end{bmatrix}$$

Fig. S1. Monthly population matrices for *Aurelia aurita* reared under low (*L*) food conditions at present water temperatures (PWT) and for a predicted winter warming trend (WW; superscript values). Each matrix comprises corrected (Appendix S1 & S2) stage-specific transition probabilities ($t$, month$^{-1}$) and fecundities ($F$, ind. ind.$^{-1}$ month$^{-1}$) for polyps (P), ephyrae (E) and medusae (M) between the months of the year: *L*1: Jan−Feb; *L*2: Feb−Mar; *L*3: Mar−Apr; *L*4: Apr−May, *L*5: May−Jun, *L*6: Jun−Jul, *L*7: Jul−Aug, *L*8: Aug−Sept, *L*9: Sept−Oct, *L*10: Oct−Nov; *L*11: Nov−Dec, *L*12: Dec−Jan. For months without survivors in the ephyra and medusa stage, the constant probability $c = 1 \times 10^{-10}$ month$^{-1}$ replaces $t_{EE} = 0$ month$^{-1}$ and $t_{MM} = 0$ month$^{-1}$ to avoid matrix reducibility.

$$A_{H1} = \begin{bmatrix} 1.010 & 0 & 0 \\ 0^{0.107} & 0.121 & 0 \\ 0^{1.377} & 0.879 & 1 \end{bmatrix}$$

$$A_{H2} = \begin{bmatrix} 1.011 & 0 & 0 \\ 0.850^{0.267} & c^{0.152} & 0 \\ 0^{2.516} & 0^{0.823} & 0.939 \end{bmatrix}$$

$$A_{H3} = \begin{bmatrix} 1.022 & 0 & 0 \\ 0.349^{0.628} & 0.152 & 0 \\ 3.394^{6.109} & 0.849 & 1 \end{bmatrix}$$

$$A_{H4} = \begin{bmatrix} 1.340 & 0 & 0 \\ 0.205^{0.370} & 0.121 & 0 \\ 2.356^{4.240} & 0.780 & 0.774 \end{bmatrix}$$

$$A_{H5} = \begin{bmatrix} 1.306 & 0 & 0 \\ 0 & c & 0 \\ 0 & 0.896 & 0.792 \end{bmatrix}$$

$$A_{H6} = \begin{bmatrix} 1.200 & 0 & 0 \\ 0 & c & 0 \\ 0 & 0 & 0.421 \end{bmatrix}$$

$$A_{H7} = \begin{bmatrix} 1.174 & 0 & 0 \\ 0 & c & 0 \\ 0 & 0 & 1 \end{bmatrix}$$

$$A_{H8} = \begin{bmatrix} 0.990 & 0 & 35481 \\ 0 & c & 0 \\ 0 & 0 & 1 \end{bmatrix}$$

$$A_{H9} = \begin{bmatrix} 0.990 & 0 & 34335 \\ 0 & c & 0 \\ 0 & 0 & 1 \end{bmatrix}$$

$$A_{H10} = \begin{bmatrix} 1 & 0 & 35477 \\ 0 & c & 0 \\ 0 & 0 & 1 \end{bmatrix}$$

$$A_{H11} = \begin{bmatrix} 1.072 & 0 & 34331 \\ 0 & c & 0 \\ 0 & 0 & 0.750 \end{bmatrix}$$

$$A_{H12} = \begin{bmatrix} 1.041 & 0 & 0 \\ 0.021^{0.038} & c & 0 \\ 0 & 0 & 1 \end{bmatrix}$$

Fig. S2. Monthly population matrices for *Aurelia aurita* reared under high (*H*) food conditions at present water temperatures (PWT) and for a predicted winter warming trend (WW; superscript values). Each matrix comprises corrected (Appendix S1 & S2) stage-specific transition probabilities ($t$, month$^{-1}$) and fecundities ($F$, ind. ind.$^{-1}$ month$^{-1}$) for polyps (P), ephyrae (E) and medusae (M) between the months of the year: *H*1: Jan−Feb; *H*2: Feb−Mar; *H*3: Mar−Apr; *H*4: Apr−May, *H*5: May−Jun, *H*6: Jun−Jul, *H*7: Jul−Aug, *H*8: Aug−Sept, *H*9: Sept−Oct, *H*10: Oct−Nov; *H*11: Nov−Dec, *H*12: Dec−Jan. For months without survivors in the ephyra stage, the constant probability $c = 1 \times 10^{-10}$ month$^{-1}$ replaces $t_{EE} = 0$ month$^{-1}$ to avoid matrix reducibility.
